# Time-resolved, deuterium-based fluxomics uncovers the hierarchy and dynamics of sugar processing by *Pseudomonas putida*

**DOI:** 10.1101/2023.05.16.541031

**Authors:** Daniel C. Volke, Nicolas Gurdo, Riccardo Milanesi, Pablo I. Nikel

## Abstract

*Pseudomonas putida*, a soil bacterium widely used for synthetic biology and metabolic engineering, processes glucose through convergent peripheral pathways that ultimately yield 6-phosphogluconate. Such a periplasmic gluconate shunt (PGS), composed by glucose and gluconate dehydrogenases, sequentially transforms glucose into gluconate and 2-ketogluconate. Although the secretion of these organic acids by *P*. *putida* has been extensively recognized, the mechanism and spatiotemporal regulation of the PGS remained elusive thus far. To address this challenge, we have developed a novel methodology for metabolic flux analysis, *D-fluxomics*, based on deuterated sugar substrates. D-Fluxomics demonstrated that the PGS underscores a highly dynamic metabolic architecture in glucose-dependent batch cultures of *P*. *putida*, characterized by hierarchical carbon uptake by the PGS throughout the cultivation. Additionally, we show that gluconate and 2-ketogluconate accumulation and consumption can be solely explained as a result of the interplay between growth rate-coupled and decoupled metabolic fluxes. As a consequence, the formation of these acids in the PGS is inversely correlated to the bacterial growth rate—unlike the widely studied overflow metabolism of *Escherichia coli* and yeast. Our findings, which underline survival strategies of soil bacteria thriving in their natural environments, open new avenues for engineering *P*. *putida* towards efficient, sugar-based bioprocesses.

## Introduction

Glycolysis, the biochemical process that converts glucose into pyruvate (Pyr, or other simple intermediates), constitutes a central part of cellular metabolism and is broadly distributed in all domains of Life [1,2]. Multiple biochemical strategies for glycolysis exist in Nature, yet our view and understanding of sugar processing are still largely dominated by the metabolic architecture found in *Escherichia coli* and *Saccharomyces cerevisiae.* In these model organisms, glucose is channeled primarily through the linear Embden-Meyerhof-Parnas (EMP) pathway towards Pyr formation [3,4]. Yet, understanding glucose catabolism in non-canonical bacterial hosts [5] is of paramount interest to engineer efficient sugar consumption pathways [6]—an aspect that has largely relied on manipulating the EMP route thus far. Indeed, the Entner-Doudoroff (ED) and the pentose phosphate (PP) pathways (or variations thereof) prevail as the main sugar catabolic route in microorganisms that live in diverse, often-changing environmental niches [7–11].

A distinct metabolic architecture ultimately leading to Pyr formation is the periplasmic gluconate shunt (PGS). *Pseudomonas* species, acetic acid bacteria and other microorganisms use these peripheral sugar oxidation routes, which generate gluconate and/or 2-ketogluconate (2KG) from glucose [7,12–14]. A peculiar feature of the PGS is that energy generation does not require active substrate uptake, unlike EMP-based sugar processing [15]. Instead, the oxidation of glucose by glucose 2-dehydrogenase (Gcd) in the PGS is directly coupled to the reduction of pyrroloquinoline quinone (PQQ) electron-carriers, while the oxidation of gluconate to 2KG is accompanied by the reduction of flavin adenine dinucleotide (FAD) [16,17]. Further oxidation of both redox cofactors during respiration yields adenosine triphosphate (ATP, with a different yield depending on the route), thus fueling the cell’s energy metabolism [18,19]. The presence of the PGS in several species underscores an ecological role for microbes thriving in their natural habitats, and gluconate excretion has been associated to gut colonialization and as a signal triggering toxin synthesis [20,21]. Some bacteria even shape the surrounding environment through gluconate secretion, as the acid helps solubilizing essential nutrients, thereby increasing their bioavailability [22,23]. Besides its ecological importance, the PGS has important implications for industrial fermentations and bioprocess engineering. *P. putida*, a soil bacterium widely adopted for metabolic engineering [24,25], favors glucose processing through the PGS [26,27]—yet the actual contribution of periplasmic oxidation reactions to the overall flux distribution has been merely estimated based on indirect experimental evidence.

The limited knowledge on these glycolytic strategies largely stems from a dearth of analytical tools to study the dynamics of periplasmic sugar processing. Metabolic flux analysis (MFA) provides valuable insight into which pathways are active and how carbon fluxes are distributed in a number of biological systems [28–30]. However, most MFA methodologies are heavily tailored for metabolic architectures found in *E. coli* and *S. cerevisiae* [31]. Adapting MFA techniques to other biological systems is a relatively straightforward task—provided that the metabolic blocks are conserved in the species of interest [32]. An interesting exception is the cyclic glycolysis of environmental bacteria, i.e. the EDEMP cycle [33], which comprises elements of the EMP, ED and PP pathways. Labeling information for metabolites derived from glucose-6-phosphate (G6P) and fructose-6-phosphate (F6P), in addition to the labeling information in proteinogenic amino acids, is needed to resolve EDEMP fluxes [34]. Similarly, the PGS presents a significant challenge to traditional MFA approaches due to several reasons. Firstly, conversion of glucose to gluconate and/or 2KG does not involve any carbon atom transition. Thus, the pattern of carbon labeling in downstream metabolites does not provide information about which pathway(s) gave rise to them. Indeed, in previous MFA studies of *P*. *putida*, the PGS fluxes could only be inferred by indirect measurement of enzyme activities *in vitro* [33–35]. Secondly, the concentrations of gluconate and 2KG are not in a *pseudo*-steady state, but they rather fluctuate during bacterial growth—leading to a substantial (yet temporary) accumulation of these organic acids in batch cultures. Therefore, a singular time-point measurement captures only a snapshot of the fluxes and generates limited information about flux dynamics over time [36].

To solve the long-standing question on the metabolic role of the PGS in *P*. *putida*, we have developed a novel MFA protocol (termed *D-fluxomics*), based on a combination of ^13^C and ^2^H-labeled substrates. ^2^H (deuterium, D) was found to be an ideal isotopic tracer to follow the fate of glucose in the upper sugar processing routes. Time-course experiments during batch cultivation on glucose and gluconate elucidated the spatiotemporal flux distribution through the different oxidation routes, in a first-case example of precise determination of PGS fluxes. The D-fluxomics approach revealed a hierarchical and highly dynamic uptake of different sugar forms, with rapid shifts between these substrates in a concentration-dependent fashion. Furthermore, we show that the temporal accumulation of gluconate and 2KG is inversely correlated to the growth rate and that this behavior can be solely explained by a model where the PGS flux is decoupled from bacterial growth. Thus, fluxes through peripheral oxidation steps decrease with faster growth—contrary to the traditional view of overflow metabolism. The impact of these findings on metabolic engineering efforts is discussed at the light of adopting sugars as the main feedstock for *Pseudomonas* fermentations.

## Experimental Procedures and Methods

### Bacterial strains, mutant construction and culture conditions

*Pseudomonas putida* KT2440 [37] and its mutant derivatives were cultivated at 30°C. For standard applications, routine cloning procedures and during genome engineering manipulations, cells were grown in LB medium (containing 10 g L^−1^ tryptone, 5 g L^−1^ yeast extract and 10 g L^−1^ NaCl, pH = 7.0) [38]. For all other experiments, *P. putida* was grown in de Bont minimal medium [39] supplemented with glucose or gluconate at the concentrations indicated in the text. If not stated otherwise, chemicals were supplied by Sigma-Aldrich Co. (St. Louis, MO, USA); [13C-U]-glucose and [2-^2^H]-glucose was purchased from Cambridge Isotope Laboratories Inc. (Tewksbury, MA, USA). *P. putida* 2KG (Δ*glk* Δ*gtsABCD* Δ*gnuK* Δ*gntT*) was created by marker-less, sequential deletion of the open reading frames of *glk*, *gtsABCD*, *gnuK* and *gntT* in wild-type strain KT2440 through homologous recombination [40,41]. Further deleting *gad* (encoding gluconate 2-dehydrogenase) in *P. putida* 2KG, using the same methodology, yielded *P. putida* Gln. Finally, *P. putida* Pgi, a strain devoid of all phosphoglucoisomerase activity, was created by marker-less elimination of *pgi-I* and *pgi-II* in *P*. *putida* KT2440. All deletions were confirmed by PCR amplification of the corresponding loci followed by DNA sequencing of the amplicons [42].

### General protocol for time-resolved D-fluxomics

For high-precision quantitative physiology and labeling experiments, *P. putida* strains were grown in 50 mL of de Bont minimal medium [39] supplemented with glucose or gluconate in a temperature-controlled, 200-mL vessel and agitated with a magnetic stirrer or in 250-mL baffled Erlenmeyer flasks as specified in the text. In all cases, the medium was inoculated with an overnight culture grown in the same conditions to an optical density measured at 600 nm (OD600) of ca 0.05 if not differently indicated. Samples were periodically withdrawn for determining OD600 values (in a 6705 UV/VIS scanning spectrophotometer; Jenway, Herlev, Denmark), concentration of sugars in the culture supernatants by HPLC [43] and to analyze the labeling pattern in relevant metabolic intermediates (as explained in the next section).

### Metabolite extraction and measurement of isotopologue composition

At selected time points, 1-mL culture aliquots were withdrawn and vacuum-filtered (Durapore^TM^ membrane filter, 0.45 μm; Sigma-Aldrich Co.). Upon filtration, the filter was re-extracted with 1 mL of acetonitrile/methanol/water [in a 40/40/20% (v/v) ratio] acidified with 0.1 M formic acid at –20°C [44]. Subsequently, cell debris were removed by centrifugation at 17,000×*g* for 2 min at 4°C. The supernatant was transferred to a new tube and solvents were removed by evaporation at 30°C for 90 min at reduced pressure (Concentrator Plus; Eppendorf SE, Hamburg, Germany). The samples were then fully dried in a freeze-dryer and stored at –80°C [45]. Prior to the analysis, the samples were reconstituted in 100 μL of HPLC-grade H2O. The metabolites were analyzed using a Prominence XR (Shimadzu Corp., Columbia, MD, USA) HPLC system coupled to a 5500 QTRAP mass spectrometer (SciEx, Framingham, MA, USA). The autosampler was cooled to 15°C. A sample of 10 μL was injected and separated in an XSelect HSS T3 150×2.1 mm^2^×2.5 μm column (Waters Corp., Milford, MA, USA). The column temperature was held at 40°C; metabolites were eluted with a constant flow rate of 0.4 mL min^−1^. Elution began with 100% buffer A [10 mM tributylamine, 10 mM acetic acid (pH = 6.86), 5% (v/v) methanol and 2% (v/v) 2-propanol]. Starting at 4 min, buffer B (2-propanol) was linearly increased to 15% at 8 min and held at 15% at 12 min. Buffer B was then linearly decreased to 0% at 13.5 min and kept at 0% until 17 min. The mass spectrometer was operated in negative mode with multiple reaction monitoring, and unit resolution for the mass filters Q1 and Q3 was used for detection and quantification. The parameters and settings used for electrospray ionization were as following: electrospray voltage, –4,500 V; temperature, 500°C; curtain gas, 40 pound square inch^−1^ (psi); CAD gas, 12 psi; gas 1 and 2, 50 psi each; and collision gas, high. Detection parameters were optimized for each metabolite, together with the expected transitions when in isotopic labelling experiments (**Table S1** in the Supplementary Material).

### Flux ratio analysis based on ^13^C- and ^2^H-isotopic tracers

A system of 15 equations, indicated as 𝑣_1_-𝑣_15_ and comprising reactions of the (incomplete) EMP, ED and PP pathways of *P*. *putida*, was used to resolve the distribution of fluxes within the PGS in *P*. *putida*. The metabolic network adopted for MFA, together with a list of the enzymes annotated to catalyze the corresponding reactions in *P*. *putida* KT2440, is shown in **Fig. 1**. The first step was calculating ϕG6P, the fraction of the G6P pool originating from glucose (abbreviated as Glu in the equations below). This parameter was calculated using either the *m*+3 or *m*+6 isotopologue fraction in labeling experiments fed with [U-^13^C]-glucose or the *m*+1 isotopologue fraction when [2-^2^H]-glucose was used as the carbon source, as indicated in **Equation 1**.

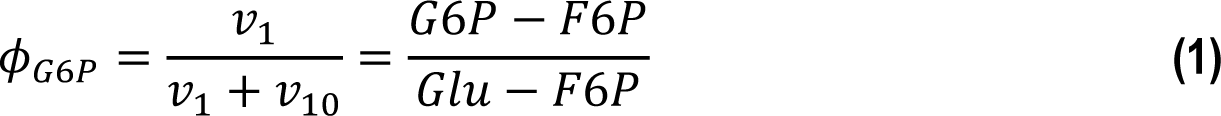

**Fig. 1.**
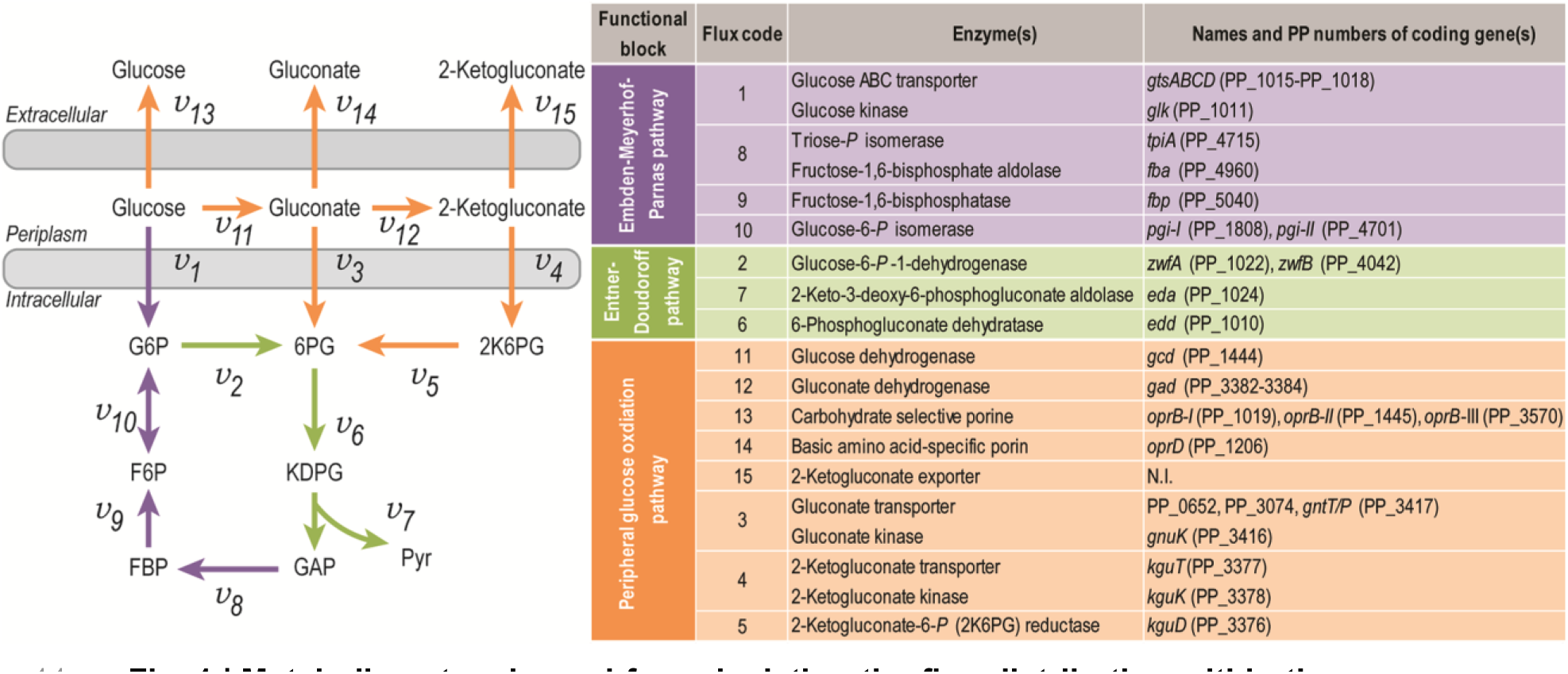
| Metabolic network used for calculating the flux distribution within the upper sugar metabolism of *P*. *putida*. The enzymes catalyzing the fluxes indicated and the genes encoding them are listed in the table; the information was compiled from the *Pseudomonas* Genome Database [92], MetaCyc [93] and the literature [94,95]. In the instances in which no gene name has been assigned, the PP number is given for each open reading frame; biochemical reactions are coded according to the three functional blocks indicated in the figure. Abbreviations: G6P, glucose-6-phosphate; F6P, fructose-6-phosphate; FBP, fructose-1,6-bisphosphate; 6PG, 6-phosphogluconate; KDPG, 2-keto-3-deoxy-6-phosphogluconate; 2K6PG, 2-keto-6-phosphogluconate; GAP, glyceraldehyde-3-phosphate; Pyr, pyruvate; *N.I.*, not identified.

It was further assumed that all flux from G6P is directed towards the formation of 6-phosphogluconate (6PG), as shown in **Equation 2**.

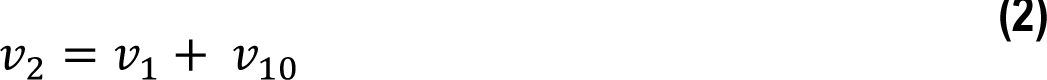

The fraction of 6PG originating form G6P (ϕ6PG) was calculated using the *m*+3 or *m*+6 isotopologue fraction in experiments with [U-^13^C] (**Equation 3**), assuming that the isotopologue fraction in glucose, gluconate (Gln) and 2KG of *m*+3 and *m*+6 is equal.

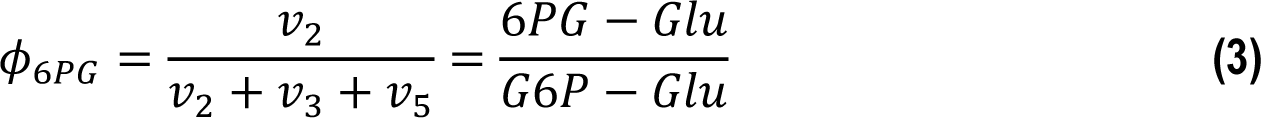

The fractional contribution of each isotopologue to the 6PG pool is given by **Equation 4**.

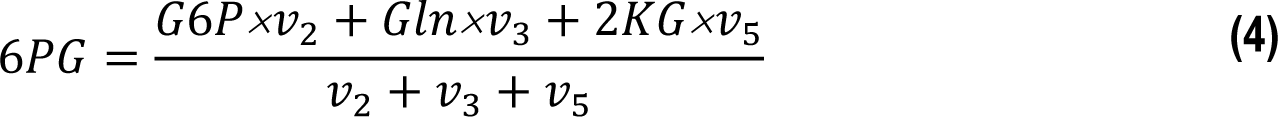

Fixing the total inflow into 6PG to 1 results in **Equation 5** and **Equation 6** for flux originating from gluconate,

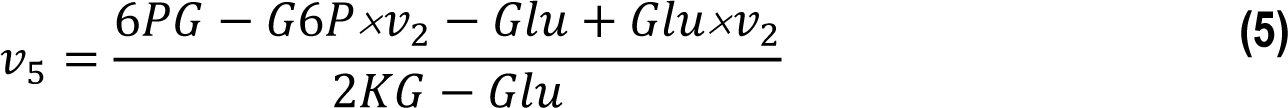

or from 2KG, *via* 2-keto-6-phosphogluconate (2K6PG).

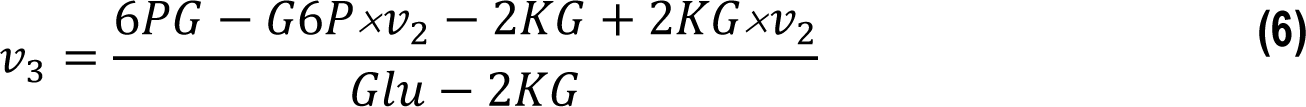

As it can be noted in the denominator of **Equations 5** and **6**, the system is solvable only if the isotopologue fraction in Glu and 2KG differ from each other. Therefore, the *m*+1 isotopologue fraction was used in experiments with [2-^2^H]-glucose as the carbon source for calculating the 𝑣_3_and 𝑣_5_fluxes.

### Quantification of the glucose, gluconate and 2-ketogluconate concentrations

An aliquot of 100-μL was withdrawn from the cultures at the indicated time-points. The samples were centrifuged at 13,000×*g* for 5 min at 4°C and the supernatant was transferred to a new tube. Subsequently, 20 μL of the supernatant were injected onto an Aminex HPX-87H (300×7.8 mm, 9 μm; Bio-Rad Laboratories Inc., Hercules, CA, USA). The sample was eluted over 30 min at 30°C with 5 mM H2SO4 at a constant flow of 0.6 mL min^−1^ [46]. Refraction index (RI) was used for the quantification of co-eluted glucose and gluconate, while absorbance at 205 nm was used to quantify gluconate and 2KG. The calculated RI of gluconate was subtracted from the detected gluconate + glucose signal to obtain the actual RI of glucose [33,47]. Authentic standards were used to construct calibration curves for all relevant sugars.

### Preparation of [2-^2^H]-labeled gluconate from deuterated glucose with resting P. putida cells

*P. putida* Gln (Δ*glk* Δ*gtsABCD* Δ*gnuK* Δ*gntT* Δ*gad*) was grown in 100 mL of de Bont minimal medium supplemented with 0.4% (w/v) succinate placed in a 500-mL baffled Erlenmeyer flask. After reaching stationary phase (ca. 18 h), cells were harvested by centrifugation at 5,000×*g* for 10 min at room temperature. The bacterial sediment was suspended in 10 mL of de Bont minimal medium without carbon source, followed by an additional centrifugation step at 5,000×*g* for 10 min. Then, cells were suspended in 50 mL of de Bont minimal media with the mixture of labeled glucose specified in the text, transferred to a 500-mL baffled Erlenmeyer flask and incubated with constant shaking at 30°C. Periodically, 1-μL aliquots were withdrawn from the culture and quickly tested for residual glucose (semi-quantitative QuantoFix^TM^ test strips; Macherey-Nagel, Düren, Germany) and pH (pH-fix 0-14 indicator strips; FischerBrand, Langenselbold, Germany). If needed, the pH was adjusted to 7.5-8 by dropwise addition of 5 M NaOH, and the transformation was continued until no residual glucose was detected. Next, the suspension was centrifuged at 5,000×*g* for 10 min, the supernatant was transferred to a new 50-mL Falcon tube (Sigma-Aldrich Co.) and centrifuged at 12,000×*g* for 10 min. Finally, the supernatant was transferred to a clean tube and filter-sterilized by passing through a 0.45-μm membrane (Sigma-Aldrich Co.). Gluconate and (residual) glucose were detected and quantified by HPLC as explained above.

### Modelling bacterial growth and extracellular metabolite concentration during batch cultivation of P. putida

The changes of metabolites in the supernatant over time are given by **Equations 7**, **8** and **9**.

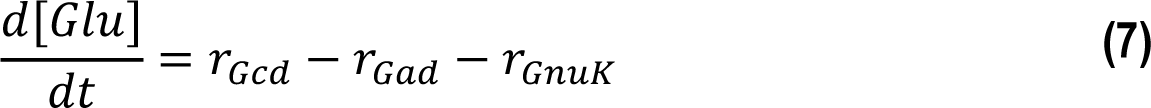

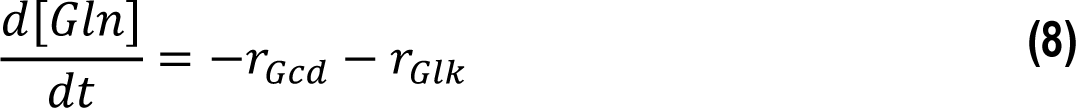

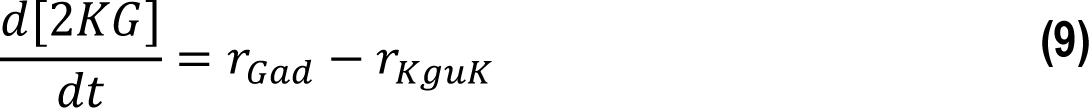

[𝐺𝑙𝑢], [𝐺𝑙𝑛] and [2𝐾𝐺] represent the concentration of glucose, gluconate and 2KG, respectively, while 𝑟_𝑍_ is the conversion rate through enzyme 𝑍. For all enzymatic steps, a Michaelis-Menten kinetic was assumed [48]. The rates through the Gad and Gcd reactions were calculated through the specific maximal activity (*V_X_^max^*) and the biomass concentration ([𝑋]) as indicated in **Equations 10 and 11**, respectively.

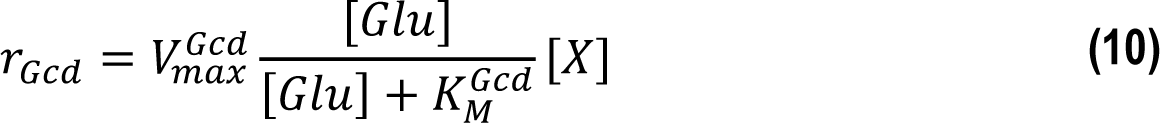

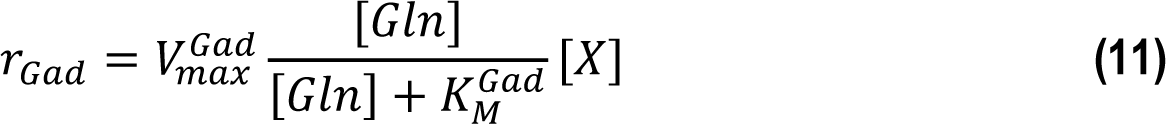

Therefore, no coupling to the growth rate (𝜇) was assumed for these calculations. In contrast, the uptake rate through the different pathways under consideration was assumed to be proportional to the growth rate, as enunciated in Equations **12**, **13** and **14**.

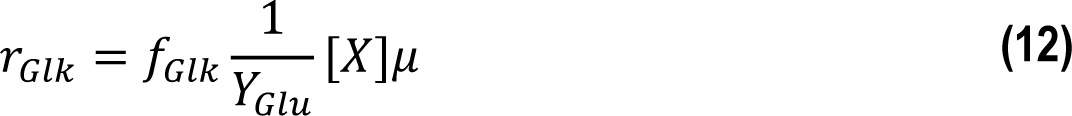

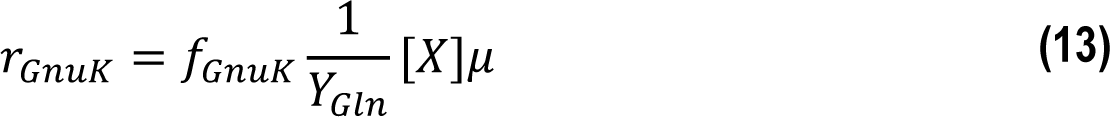

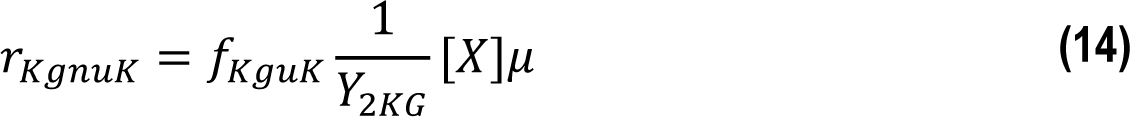

where 𝑌_𝑍_ is the yield on substrate 𝑍 and 𝑓_𝑍_ is the relative fraction of total carbon source entering through the enzymatic step 𝑍. The fractional uptake was given by the empirically-determined uptake distribution 𝐷 under saturation conditions normalized to the relative saturation of each uptake with its substrate 𝑆 (**Equation 15**).

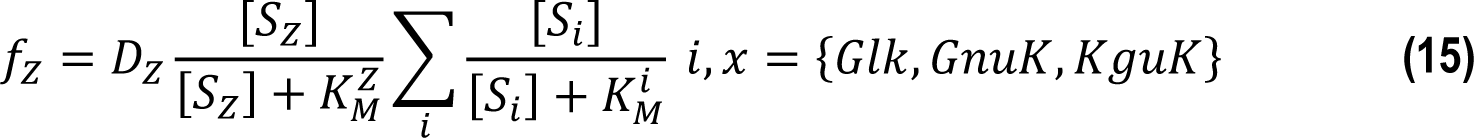

The biomass concentration was determined by **Equation 16**.

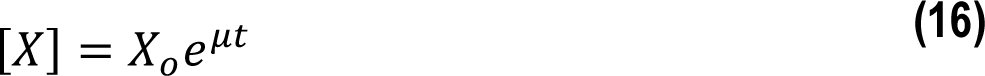

where 𝑋_0_ is the initial biomass.

Simulations were carried out with different growth rates, initial values of biomass, glucose concentration and altering kinetic parameters as indicated in the text. In all cases, at least 2,000 iterative steps were carried out with time steps between 0.001 and 0.0035 h.

### Phylogeny distribution of the periplasmic gluconate shunt across the tree of Life

Information on orthologues to Gcd (K00117; EC 1.1.5.2) and Gad (K05308; EC 4.2.1.140) was retrieved from KEGG [49] through AnnoTree [50]. The minimum identity for the search was set to 30%, while subject and query alignment were fixed to 70 and maximum *E* value to 0.00001, respectively. The resulting phylogenetic tree was drawn with the *i*TOL package [51].

### Data and statistical analysis

All experiments reported in this study were independently repeated at least twice (as indicated in the corresponding figure legend), and the mean value of the corresponding parameter ± standard deviation is presented. Whenever relevant, the level of significance of the differences when comparing results was evaluated by means of the Student’s *t* test with α = 0.01 or α = 0.05 as indicated in the figure legends.

## Results and Discussion

### The periplasmic gluconate shunt, widely distributed in the bacterial kingdom, underscores energy and redox versatility

The PGS reactions have been recognized as the predominant sugar processing mechanism in *Pseudomonas* and related rhizobacteria a long time ago [26]—yet the extent of its presence in Nature remains unsolved. To gain insight into the distribution of the PGS across multiple species in the bacterial kingdom, we selected the Gcd and Gad enzymes as a proxy of the presence of these two oxidation steps. The resulting phylogenetic reconstruction indicated that the PGS is indeed widely distributed in bacterial species (**Fig. 2A**). The PPQ-dependent glucose dehydrogenase (Gcd) was found in 31% of all bacterial species, with an uneven distribution over the different phyla (i.e. Actinobacteria, Firmicutes, Patescibacteria, Bacteroidota and Proteobacteria). In particular, the Actinobacteriota and Proteobacteria phyla were especially enriched in species harboring the PGS (**Fig. 2B**). Inside the Proteobacteria phylum (and out of the 9,474 members analyzed herein), the Pseudomonadales and Rhizobiales orders had the highest density of Gcd-bearing species—as it was expected based on the available experimental evidence. Indeed, the fraction of Gcd-containing species accounted for >0.45 of all γ-Proteobacteria (i.e. Pseudomonadales and Rhizobiales) and Actinobacteria analyzed. In contrast, Cyanobacteria had the lowest abundance of Gcd-bearing species (**Fig. 2B**). The second step in the PGS, catalyzed by gluconate dehydrogenase, was present in ca. 8% of all bacterial genomes used in this analysis, roughly following the same distribution trend as observed for Gcd—albeit at a significantly lower frequency (details not shown). The uneven distribution of the PGS in the bacterial kingdom may be connected to some intrinsic characteristics of such a metabolic feature. Firstly, gluconate cannot be metabolized through the linear EMP pathway. Thus, bacteria exclusively adopting the EMP pathway for glycolysis do not usually have an active PGS metabolism—with *E*. *coli* as a prime example [52]. Secondly, the synthesis of the PQQ cofactor is strictly oxygen-dependent [53], which indicates that the presence of a PQQ biosynthesis pathway is largely restricted to either aerobic microorganisms or those dwelling in environments where PQQ is available [54]. Interestingly, the abundance of the PGS feature in bacteria correlates well with the phylogenetic distribution of the ED catabolic route [10,55] and the presence of a PQQ biosynthesis pathway [54] over all the phyla.

**Fig. 2.**
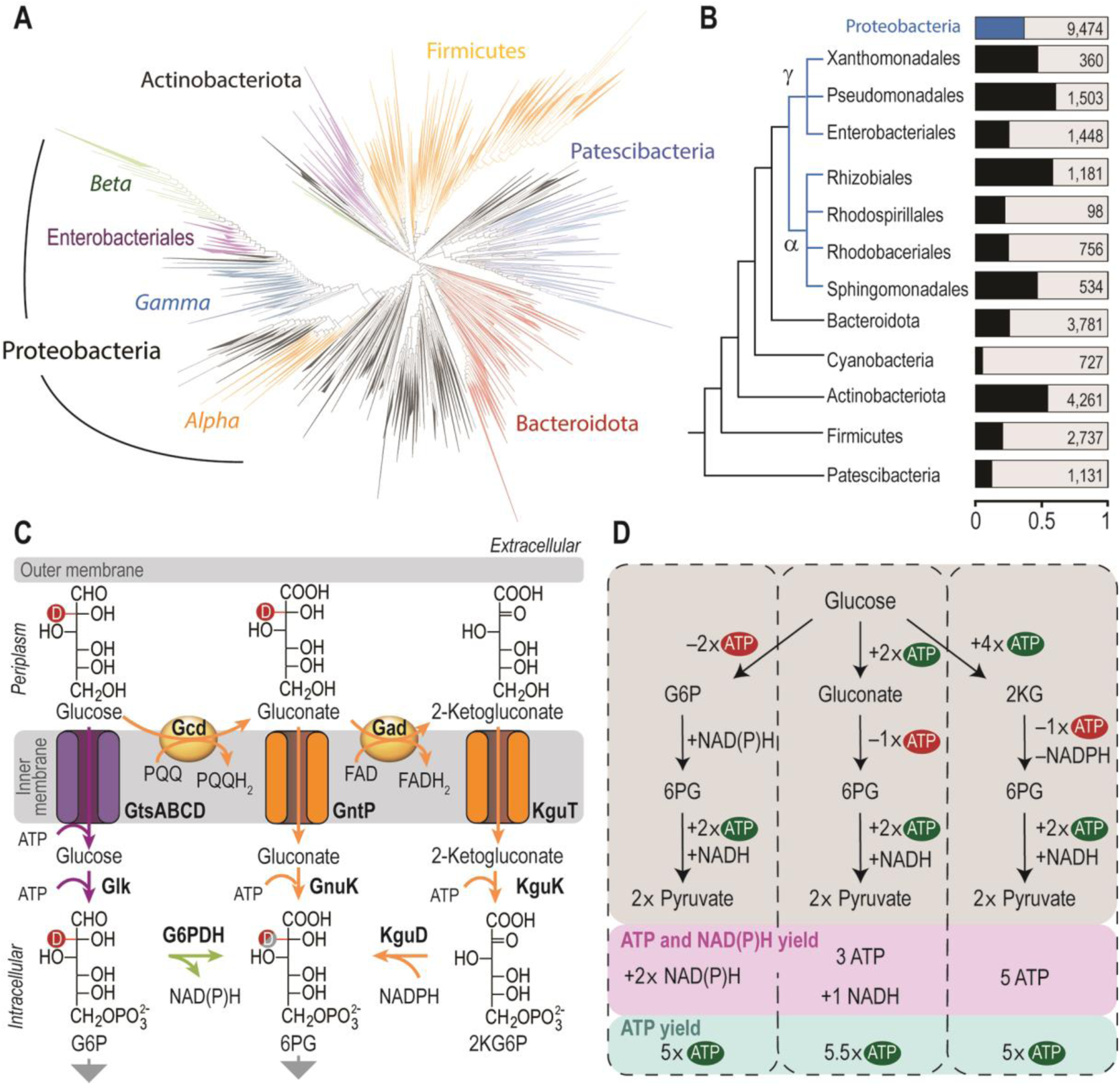
| Phylogenetic distribution and stoichiometric analysis of the periplasmic oxidation shunt. **(A)** Phylogenetic distribution of a periplasmic gluconate shunt in the bacterial kingdom. The presence of a pyrroloquinoline quinone (PQQ)-dependent glucose dehydrogenase (Gcd) was adopted as a proxy for the analysis, with a wide distribution over all phyla (overall, 31% of all bacterial species carry a Gcd polypeptide). **(B)** Main phyla and orders of bacteria harboring Gcd. Both the relative fraction of Gcd-containing bacteria and the total number of species analyzed in each phyla or order are indicated in the diagram; Proteobacteria (α and γ) are highlighted in blue. **(C)** Glucose processing through either its direct phosphorylation or periplasmic, sequential oxidation in *P*. *putida*. Glucose can be directly taken up and phosphorylated to glucose-6-phosphate (G6P) by an ATP-dependent glucose transporter (GtsABCD) and glucose kinase (Glk), respectively. The hexose can be also oxidized to gluconate *via* glucose dehydrogenase (Gcd). Similarly, gluconate can be transported by a gluconate permease (GntP) and either phosphorylated by gluconokinase (GnuK) to 6-phosphogluconate (6PG) or oxidized to 2-ketogluconate through a flavin adenine dinucleotide (FAD)-dependent gluconate dehydrogenase (Gad). Finally, 2-ketogluconate is imported to the bacterial cytoplasm and phosphorylated to 2-keto-6-phosphogluconate (2K6PG) by means of the 2-ketogluconate transporter (KguT) and a 2-ketogluconate kinase (KguK), respectively. Note that deuterium (D, indicated as a red circle) at position two in the glucose molecule would remain unaffected during glucose and gluconate uptake; however, gluconate oxidation to 2-ketogluconate releases the deuterium label (see also Fig. 3). Therefore, the 6PG pool will be differentially labeled depending on which sugar uptake routes are active. Grey arrows indicate further processing of key intermediates into central carbon metabolism. **(D)** Formation and consumption of ATP and soluble reducing equivalents [i.e. NAD(P)H] through the different catabolic routes of glucose to pyruvate. The ATP yield *via* Gad processing was estimated based on the translocation of 4 H^+^ [96].

To understand the metabolic function of the PGS, it is worth to analyze the stoichiometry and the energy and redox yields of these routes in detail. The PGS directly couples the oxidation of glucose and gluconate to the respiratory chain. In particular, the oxidation of glucose to glucono-δ-lactone, an intermediate that spontaneously hydrolyses to gluconate, drives PQQ reduction to PPQH2, while the oxidation of gluconate to 2KG is coupled to FAD reduction to FADH2 (**Fig. 2C**). Both reduced cofactors subsequently transfer the electrons to ubiquinol, which allows for the direct translocation of two protons [56]. In contrast, the oxidation of ubiquinone, the reduced form of ubiquinol, enables the translocation of six protons through the complexes III and IV in the respiratory chain. Taken together, these steps yield eight translocated protons—while the oxidation of NAD(P)H, the redox currency formed in the alternative, cytoplasmic and ATP-dependent sugar oxidation, results in the translocation of ten protons through complexes I, III and IV in the respiratory chain. Assuming that four protons drive the synthesis of one ATP molecule, the metabolic route that proceeds through G6P dehydrogenase (G6PDH) and 2K6PG reductase (KguD) yields 5 ATP equivalents, while the direct phosphorylation of gluconate yields 5.5 ATP (**Fig. 2D**). Even though the ATP yields are in principle similar, the balance of redox cofactors in the cell differs significantly. While the uptake of glucose and further processing to 6PG produces NAD(P)H [55], 2KG transport and conversion to 6PG consumes redox equivalents [14,57]—while gluconate uptake (and direct phosphorylation to 6PG) is redox-neutral [58]. Such differences at the energy and redox balances probably confer metabolic flexibility to bacteria running these alternative glycolytic strategies in parallel [59–61]— without any significant protein expenditure. Motivated by these observations on the ubiquitous presence of the energy and redox versatile PGS-based metabolism in environmental bacteria, we adopted *P*. *putida* as a model system to explore the mechanistic details of glucose transport, oxidation and downstream catabolism as explained in the next section.

### D-Fluxomics: adopting [2-^2^H]-glucose as an isotopic tracer for metabolic flux analysis

MFA methodologies have been adapted to analyze a broad range of organisms beyond *E. coli*, yeast and mammalian cells [31]. As indicated previously, novel challenges had to be solved prior to applying ^13^C-based MFA in *P*. *putida* and related species. While glucose is metabolized through a linear EMP pathway in *E*. *coli* and yeast, rhizobacteria harbor a cyclic glycolytic catabolism [33,59]. In addition to this occurrence, the distribution of fluxes in the PGS cannot be fully elucidated with the currently available MFA methods [34]—owing to identical carbon transitions in the two branches of the PGS, which preclude labeling-based discrimination. To address this technical challenge, we evaluated ^2^H-based MFA as an alternative methodology to elucidate fluxes through the PGS node. We started by evaluating the atomic transitions expected in the PGS if *P*. *putida* KT2440 was grown on [2-^2^H]-glucose (**Fig. 2C**). The D label at position C2 of the sugar should be conserved in the reactions catalyzed by Glk, Gcd and GnuK, while the oxidation of gluconate to 2KG by Gad and its subsequent reduction by KguD would lead to the replacement of the D label by an H atom derived from water or NADPH (**Fig. 2C**). Therefore, the enrichment in deuterated 6PG relative to G6P should reflect the flux ratio through GnuK and KguK. Moreover, the pattern of D-labeling of F6P provides useful information on which glycolytic pathway gave rise to this metabolite (**Fig. 3A**). Indeed, the conversion of G6P into F6P by Pgi (phosphoglucoisomerase) does not affect the labeling in the C2 position, whereas D is lost when G6P is processed through the EDEMP cycle. When F6P is derived from 6PG through the PP pathway, the resulting labeling pattern corresponds to a mixed pool of D-containing and D-free F6P molecules. If the information based on D enrichment of these metabolites is combined with an orthogonal isotopic tracer, e.g. [U-^13^C]-glucose, this methodology, termed *D-fluxomics*, should enable resolving all fluxes of carbon uptake and cycling through the EDEMP cycle.

**Fig. 3.**
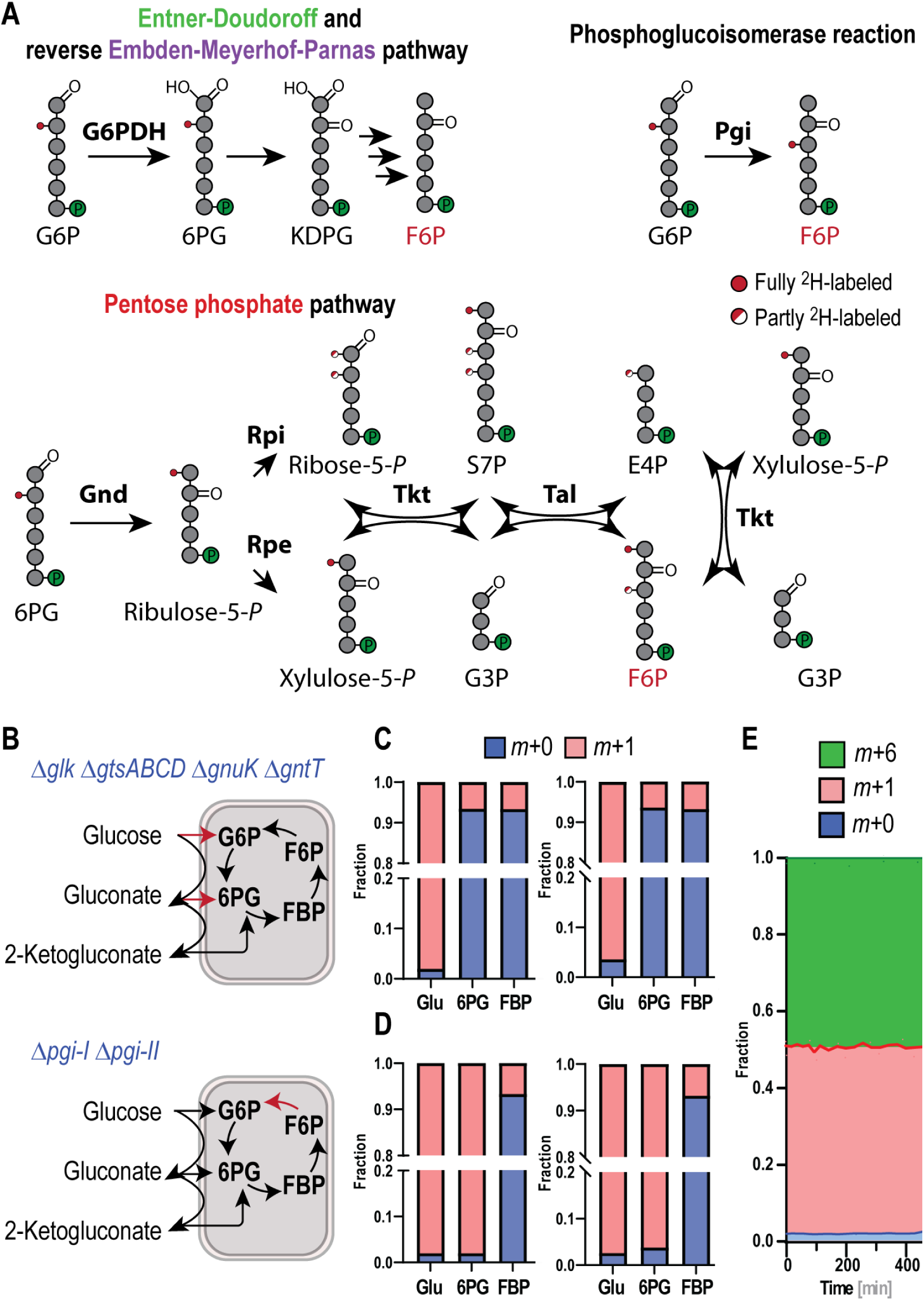
| Deuterium transition maps and experimental validation of D-fluxomics in *P*. *putida* grown on [2-^2^H]-glucose. **(A)** Atomic transitions through the Entner-Doudoroff (ED) pathway, the reverse (i.e. gluconeogenic) Embden-Meyerhof-Parnas (EMP) pathway and the pentose phosphate (PP) pathway. The deuterium (D, ^2^H) label is lost in the ED catabolism during the oxidation of 6-phosphogluconate (6PG) to 2-keto-3-deoxy-6-phosphogluconate (KDPG). In the EMP pathway, [2-^2^H]-glucose-6-phosphate (G6P) can be converted into [3-^2^H]-fructose-6-phosphate (F6P) by the action of phosphoglucoisomerase (G6P isomerase, Pgi). In contrast, labeling transitions in intermediates of the PP pathway are more complex and enzyme-dependent. Glyceraldehyde-3-phosphate (G3P) remains unlabeled regardless of the enzyme activities involved in 6PG processing, while F6P is D-labeled in position C1 and, depending on its metabolic origin, also in position C3. In all cases, the D signal is marked with a red dot; if the probability of finding a D label in the annotated position is 50%, the dot is half filled. Abbreviations: G6PDH, G6P dehydrogenase; Gnd, 6PG dehydrogenase; Rpe, ribulose-5-phosphate 3-epimerase; Rpi, ribulose-5-phosphate isomerase; Tkt, transketolase; Tal, transaldolase; S7P, sedoheptulose-7-phosphate; and E4P, erythrose-4-phosphate. **(B)** Strains used for experimental validation of D-fluxomics. *P. putida* 2KG (Δ*glk* Δ*gtsABCD* Δ*gnuK* Δ*gntT*) cannot uptake glucose or gluconate, but this strain is still able to oxidize glucose to 2-ketogluconate, which is transported and processed to 6PG (upper panel). Deletion of the two phosphoglucoisomerase genes of *P*. *putida* (i.e. *pgi-I* and *pgi-II*) interrupts carbon cycling in the EDEMP cycle (lower panel). The missing reactions in the mutant *P*. *putida* strains are indicated with red arrows. **(C)** Theoretical and experimentally-determined D-label patterns (left and right panel, respectively) for glucose (Glu), 6PG and fructose-1,6-bisphosphate (FBP) when *P. putida* 2KG was grown on 100% [2-^2^H]-glucose. The molecular weight of these metabolites is represented as a fraction of the total pool for both the native form devoid of any heavy isotope (*m*+0) and molecules containing a single D isotope (*m*+1). **(D)** Theoretical and experimentally-determined D-label patterns (left and right panel, respectively) for glucose (Glu), 6PG and fructose-1,6-bisphosphate (FBP) calculated and determined when *P*. *putida* Pgi was grown on 100% [2-^2^H]-glucose. **(E)** Fractional labelling of glucose throughout a batch fermentation of wild-type *P*. *putida* KT2440. In these experiments, the initial isotope tracer composition was 50% [U-^13^C]-glucose and 50% [2-^2^H]-glucose. The *m*+0, *m*+1 and *m*+6 fractions of the glucose pool are individually indicated; all experiments represented in the figure were repeated in independent biological triplicates.

In order to explore the metabolic H/D transitions *in vivo* and validating the D-fluxomics methodology, we constructed two mutants of *P. putida* KT2440, termed strains 2KG and Pgi (**Fig. 3B**). Owing to the deletions in *glk*, *gtsABCD*, *gnuK* and *gntT*, glucose can be only oxidized to gluconate or 2KG by *P*. *putida* 2KG—but only 2KG, the most oxidized form of the sugar, can be transported into the cell and reduced to 6PG. In contrast, elimination of *pgi-I* and *pgi-II* (giving rise to *P*. *putida* Pgi) blocked the interconversion of G6P and F6P—effectively interrupting cycling of hexoses phosphate within the EDEMP cycle. These strains were grown in shaken-flasks cultures using [2-^2^H]-glucose as the only carbon source, and the labeling pattern of glucose, 6PG and FBP (i.e. the *m*+0 and *m*+1 pools for each metabolite) was determined during exponential growth (**Fig. 3C** and **D**). As predicted from the atomic transitions within the PGS (**Fig. 2C**), the D labeling in position C2 of glucose should be lost during carbon uptake *via* 2KG in *P*. *putida* 2KG—and the intermediate pools should reflect the natural isotope abundance of glucose. Indeed, growing the mutant strain 2KG on 100% [2-^2^H]-glucose led to the expected labeling pattern in 6PG and FBP with a high level of accuracy (with < 8% of the hexose phosphate pool containing a D label at position C2; **Fig. 3C**). Hence, these experimental results confirm the proposed H/D transitions through the sequential activities of Gad and KguK (**Fig. 2C**). In order to check if the D-label is lost during carbon uptake at the level of glucose or gluconate (and their processing through Glk and GnuK), *P. putida* Pgi was likewise grown on [2-^2^H]-glucose. As indicated above, knocking-out the phosphoglucoisomerase step prevents cycling through the EDEMP pathway, which would otherwise result in an increased fraction of unlabeled G6P and 6PG as the D in position C2 is eliminated in the dehydration of 6PG by Edd. Again, the experimentally-determined labeling pattern of metabolites extracted from *P*. *putida* Pgi matched the theoretical prediction almost identically (**Fig. 3D**). In this case, the labeling of the G6P pool mirrored the labeling pattern in glucose, while the minor fraction of deuterated FBP could only be attributed to the natural isotopic abundance in the substrate.

While these results indicated the general validity of the D-fluxomics approach, a last concern was the stability and persistence of the [2-^2^H]-glucose–dependent labeling in the metabolites, as spontaneous proton exchange reactions exist between carbohydrates and water [62,63]. To test whether this phenomenon could affect our experimental setup, a mixture of 50% [U-^13^C]-glucose and 50% [2-^2^H]-glucose was used as the only carbon substrate in shaken-flask cultivations of wild-type *P. putida* KT2440, and the isotopologue composition of the residual glucose was periodically monitored throughout the batch cultivation (**Fig. 3E**). We detected no noticeable changes in the isotopologue composition of glucose over time. This observation suggest that, on the one hand, the D label is stable during the timescale relevant for D-fluxomic experiments and, on the other hand, that there is no measurable discrimination of isotopologues by the enzymes involved in sugar uptake and oxidation in *P*. *putida*. Even when a slight preference against isotopes by some enzymes cannot be totally ruled out, such an effect has been deemed negligible for most tracer experiments [64,65]. The labeling strategy presented in this section satisfies all of the requirements needed to be adopted for metabolic flux analysis—hence, [2-^2^H]-glucose was employed for elucidating sugar metabolism in *P. putida* as disclosed below.

### Deploying D-fluxomics for high-resolution mapping of glucose processing in P. putida

#### Challenges in metabolic flux analysis of non-canonical sugar processing architectures

Thus far, MFA studies on *P. putida* and related species have been carried out essentially at single-time point measurements during exponential growth [33-35,59,66,67]. As indicated above, such procedures were adopted from methodologies that were originally designed for *E. coli* and yeast cultivations. In these model organisms, all metabolites reach a *quasi*-steady-state during exponential growth, except for the substrate, glucose, and secretion products, e.g. acetate, ethanol and CO2 [68]. Under these conditions, the intracellular metabolite concentration (hence, the producing and consuming fluxes) does not change over time. This assumption, however, does not hold true for *Pseudomonas*. In these species, gluconate and 2KG accumulate in the culture medium throughout the cultivation on glucose, which indicates that the producing and consuming fluxes around these metabolites are not constant. From a methodological point of view, this problem is compounded by the dearth of experimental data on the enrichment of gluconate and 2KG in the bacterial periplasm. Furthermore, the extensive lag phases observed in cultures of *P*. *putida* mutants defective in direct glucose utilization in the EDEMP cycle (Δ*zwfA* Δ*zwfB* Δ*zwfC*) or glucose and gluconate processing *via* phosphorylation (Δ*glk* and Δ*gnuK*, respectively) strongly suggest that *pseudo*-steady state concentrations for gluconate and 2KG are not immediately established in the periplasm [55]. Furthermore, the primary literature reports significant differences in the flux distribution between sugar phosphorylation or oxidation. Such discrepancies range from 10% to 33% for direct glucose phosphorylation, while other fluxes vary much less between studies [33,34]. Taken together, the observations listed above gave rise to the hypothesis that the initial steps of glucose routing and processing in *P*. *putida* are dynamic, and probably dependent on the growth phase and oxygenation of the culture. Therefore, we aimed at capturing flux changes during batch cultivations by performing time-series measurements instead of single sampling points and integrating the data in the context of the D-fluxomics approach. Here, measuring the labeling pattern of glycolytic intermediates (instead of the customary protein-derived amino acids) is advantageous as the half-life time of these metabolites is much shorter (in the order of seconds to minutes)—therefore, they retain information on the metabolic fate of carbon atoms during short periods [69,70]. The labeling patterns of proteinogenic amino acids, in contrast, convey information of a longer period of biochemical processing. Building on these considerations, we adopted the D-fluxomics framework to obtain a detailed map of the glucose processing routes in *P*. *putida*.

#### Hierarchical uptake of glucose, gluconate and 2-ketogluconate by P. putida KT2440

In order to combine experimental data on quantitative physiology, biomass formation, accumulation of metabolites in culture supernatants, together with the actual flux distribution in glucose-grown *P*. *putida*, we followed the workflow outlined in **Fig. 4A**. To this end, wild-type *P. putida* KT2440 was cultured in de Bont minimal medium containing U-^13^C, 2-^2^H and unlabeled glucose; measurements were periodically taken in parallel for all relevant parameters. The starting cell density for these cultivations was set to an OD600 of 0.2. This fairly high amount of cells at the onset of the cultivation was deliberately chosen to ensure that sufficient biomass for metabolite extraction procedures would be available shortly after initiating the cultures—thereby capturing the metabolism in a state where glucose should be predominantly taken up and phosphorylated, rather than being oxidized to gluconate. The growth rate over the whole cultivation was essentially constant at μ = 0.68 ± 0.05 h^−1^, the biomass yield on the total carbon substrate (i.e. glucose, gluconate and 2KG) was *Y*X/S = 0.110 ± 0.004 gCDW mmol^−1^ and the total carbon uptake rate was 6.2 ± 0.5 mmol gCDW^−1^ h^−1^ (**Fig. S1A** and **S1B** in the Supplementary Material), similar to previously reported values [33,34]. The first sampling was carried out after 9.6 min of initiating the cultivation and, even at this early time point, gluconate and 2KG were already present in the extracellular medium at 0.73 ± 0.05 mM and 0.18 ± 0.01 mM, respectively (**Fig. 4B**). This observation implies an extremely fast accumulation of these sugar metabolites in the early growth phase. Even though gluconate and 2KG were present at substantial, non-negligible quantities at this data point, the flux through Glk was more than twice as high (17% ± 4%) as compared to all of the later time points (6%-9%, **Fig. 4C**)—indicating a dynamic shift from direct glucose uptake to gluconate processing. These figures are in accordance with previous fluxomics studies, where the flux through Glk ranged from 10%-33% of the total carbon uptake. Importantly, the observed flux variation over time (**Fig. 4C**) contradicts the prevailing hypothesis that gluconate and 2KG reach a *pseudo*-steady state in the bacterial periplasm. Hence, a gradual transition from pure glucose phosphorylation towards a mixed sugar uptake seems to occur during the early phase of cultivation. While the difference between the flux through Glk at the first to all other time points was statistically significant (*P* < 0.05, Student’s *t*-test), there was no relevant difference between the later time points. It seems plausible that glucose uptake and processing through Glk remains constant over time (at an average flux of 7.3% ± 1.5%) once sufficient amounts of gluconate and 2KG have accumulated in the medium. Indeed, the Glk flux was maintained within this range even when the glucose concentration dropped below the limit of quantification (∼0.1 mM) at 350 min (**Fig. 4B** and **4C**)—most likely due to the high-affinity nature of glucose uptake in *P*. *putida*. Orthologues of the glucose import system in other *Pseudomonas* species exhibit *K*M values in the low-μM range (e.g. *P. aeruginosa*, *P. chlororaphis* and *P. fluorescens*; **Table S2** in the Supplementary Material), thus supporting this notion.

**Fig. 4.**
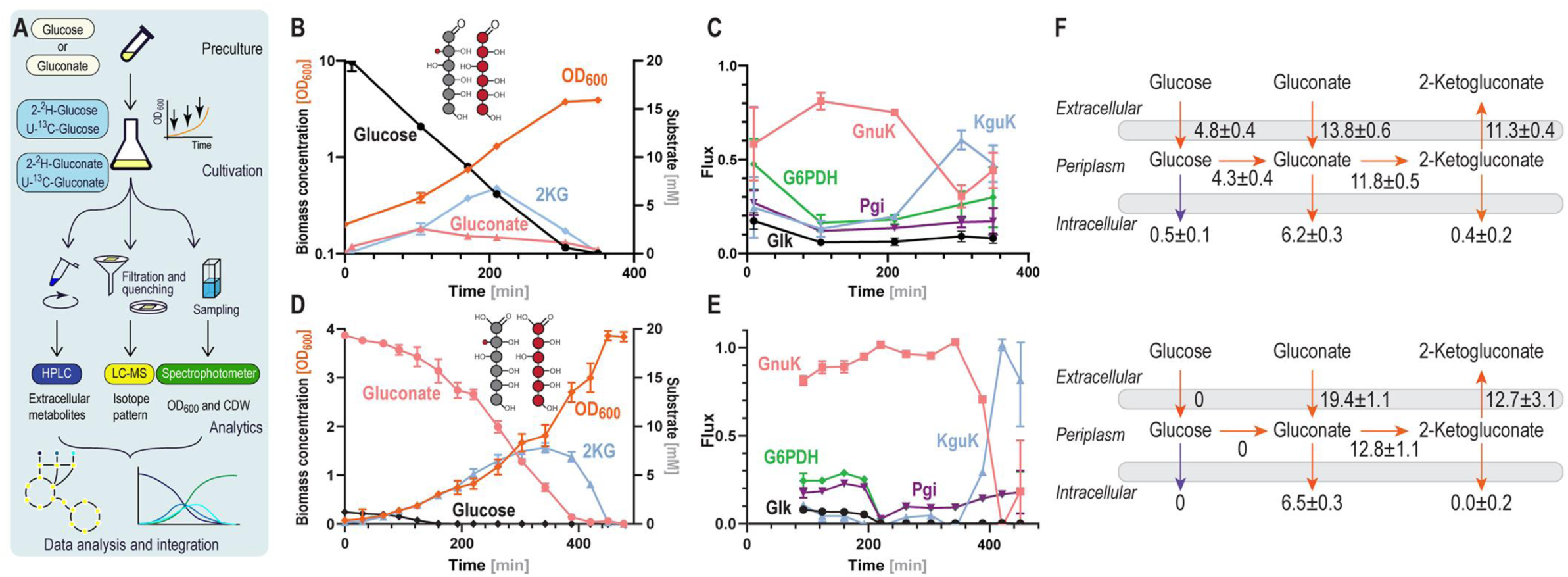
| Time-resolved D-fluxomic analysis for *P*. *putida* KT2440 growing on glucose or gluconate. **(A)** Workflow for time-resolved D-fluxomics. Bacteria were cultivated in de Bont minimal medium supplemented with labeled (either ^13^C or ^2^H) glucose. Over the cultivation, samples were periodically taken to determine cell density (optical density at 600 nm, OD600, and cell dry weight, CDW), concentration of carbohydrates in the supernatant (by HPLC) and the labeling pattern of key metabolites (by LC-MS). These data were then combined to identify metabolic routes active during glucose-dependent batch cultivations. **(B)** Biomass and carbohydrate concentration profiles in batch cultures of wild-type *P*. *putida* KT2440 grown on glucose. Cells were grown on a mixture of 25% unlabeled glucose, 25% [2-^2^H]-glucose and 50% [U-^13^C]-gluconate (total carbohydrate concentration = 20 mM). **(C)** Fluxes in the upper metabolism of *P*. *putida* KT2440 over time in glucose cultures. All fluxes were normalized to the total carbon uptake (i.e. *qS* = 1). Abbreviations: Glk, glucose kinase; GnuK, gluconokinase; Pgi, phosphoglucoisomerase; G6PDH, glucose-6-phosphate dehydrogenase; KguD, 2-ketogluconate-6-phosphate reductase. Data represent mean values ± standard deviation of three independent experiments. **(D)** Biomass and carbohydrate concentration profiles in batch cultures of wild-type *P*. *putida* KT2440 grown on gluconate. Cells were grown on a mixture of 50% [U-^13^C]-gluconate and 50% [2-^2^H]-gluconate (total carbohydrate concentration = 20 mM; note that ca. 1 mM residual glucose was present in the medium at the onset of the cultivation). **(E)** Fluxes in the upper metabolism of *P*. *putida* KT2440 over time in gluconate cultures. All fluxes were normalized to the total carbon uptake (i.e. *qS* = 1); data represent mean values ± standard deviations of three experiments. **(F)** Sugar uptake and distribution of fluxes within the periplasmic glucose oxidation in *P*. *putida* KT2440. Fluxes are shown for the early growth phase (upper panel) and the late growth phase, after gluconate depletion (lower panel). Fluxes were calculated from the relative uptake rates and biomass yields on the different substrates. In this metabolic map, fluxes are given as mmol gCDW^−1^ h^−1^; the growth rate in the early growth phase was μ = 0.78 ± 0.08 h^−1^ and μ = 0.38 ± 0.04 h^−1^ in the late growth phase.

Gluconate processing through GnuK dominated the overall carbon uptake profile during most of the experiment—except for the very late stage of the cultivation, when the gluconate concentration fell below 1.12 ± 0.08 mM (**Fig. 4B**). The main carbon uptake regime, in the form of gluconate, is fully aligned with previous reports [33–35], where early-exponential phase cultures (OD600 = 0.2-0.6) were used to determine fluxes at a single time point. However, our data indicates that a significant fraction of 2KG was also taken up while there was more than 1 mM gluconate still present in the culture medium. An early study reported a *K*M value for GnuK = 0.12 mM [71], while the *K*M of the gluconate transport system has been determined to be ∼0.5 mM [58]. These kinetic parameters determined *in vitro* help explaining our results, as the gluconate importer would not be operating at saturation, which is directly linked to the decreasing flux through this route over time. Uptake, phosphorylation and reduction of 2KG through the sequential action of KgnuT, KguK and KguD remained low (<20%) until the late fermentation phase. A significantly increased 2KG uptake (60% ± 5%) occurred only when the gluconate concentration had dropped below a threshold concentration while 2KG was still present at relatively high amounts (at *t* = 305 min). Furthermore, when all available carbon substrates (i.e. glucose, gluconate and 2KG) were nearly depleted (at *t* = 350 min), the magnitude of the fluxes through GnuK and KguK was in a similar range (44% ± 9% and 48% ± 9%, respectively; **Fig. 4C**). A reduction in the specific growth rate was observed at this phase—probably due to the fact that the overall carbon uptake rate was insufficient to support unlimited bacterial growth.

While the fluxes through the convergent uptake pathways displayed high plasticity during the whole batch cultivation, the fluxes within the EDEMP cycle remained fairly stable over time—as represented by the Pgi flux (**Fig. 4C**). Indeed, the Pgi flux was significantly elevated (27 ± 7%) only at early time points (*t* < 20 min), whereas no major differences were detected between the samples taken at later time points (12%-17%). G6PDH followed a similar trend as described for the Pgi activity: a high flux was detected at the first time point (48% ± 13%), with a relatively stable value during late cultivation (16%-30%). The highest Pgi and G6PDH fluxes in the first experimental measurements may be directly connected to the larger flux through Glk. By the end of the cultivation, a significantly higher flux through KguK, the phosphorylation step prior to the activity of KguD, a strictly NADPH-dependent dehydrogenase [33], was accompanied by a slight increase in the EDEMP cycle fluxes—as indicated for the G6PDH flux. This observation may underlie a compensatory phenomenon, as there are only a few NADPH-producing reactions in the central carbon metabolism of *P. putida*, i.e. G6PDH, 6PGDH, MaeB, ICTDH and GDH [72]. Moreover, these results are in line with the general defense mechanism deployed when *P. putida* deals with oxidative stress (and the associated NADPH demand), i.e. activation of the oxidative branch in the PP pathway and increased activities through G6PDH, caused by higher flux through Glk and recycling through Pgi [35].

#### High-definition, time-resolved flux analysis of gluconate metabolism in P. putida KT2440

The dynamic changes in substrate consumption motivated us to conduct experiments at an even higher level of resolution, as the transition from gluconate-to-2KG uptake in glucose cultures happened rapidly (**Fig. 4C**). To this end, we slightly changed the experimental setup by using a mixture of 50% [U-^13^C]- and 50% [2-^2^H]-gluconate as the carbon source instead of glucose. We reasoned that this strategy would allow for enhanced and faster 2KG accumulation, thus enabling a better resolution of carbon uptake dynamics at late stages of the cultivation. Since [2-^2^H]-gluconate is not commercially available, we prepared this labeled carbon substrate in a biotransformation reaction with resting cells of *P. putida* Gln (Δ*glk* Δ*gtsABCD* Δ*gnuK* Δ*gntT* Δ*gad*) incubated in the presence of [2-^2^H]-glucose. Under our experimental conditions, resting-cell biotransformation experiments typically proceeded with a >95% efficiency in glucose oxidation. Also, in previous experiments carried out as shaken-flask cultivations, the cultures were shortly interrupted (< 1 min) for sample withdrawal. To increase the resolution and quality of our experimental data, a frequent sampling routine was implemented by carrying out *P*. *putida* cultivations in temperature-controlled, baffled vessels with aeration provided by continuous magnetic stirring. This mini-bioreactor setup facilitated frequent sampling without significantly altering the culture conditions, and preliminary experiments showed that the overall physiology of *P*. *putida* KT2440 (e.g. specific growth rate and final OD600 values) was not significantly modified under these culture conditions as compared with shaken-flask cultivations (data not shown).

The novel cultivation setup led to the formation of high amounts of 2KG by *P*. *putida* (**Fig. 4D**), with up to 7.8 ± 0.5 mM 2KG accumulated after 343 min. The cultivation started with 1.2 ± 0.2 mM [2-^2^H]-glucose still remaining in the culture medium, which dropped under the detection limit after 180 min. Interestingly, the Glk flux remained at relatively constant levels (7.2% ± 0.7%) while glucose was still detected (0.028 ± 0.002 mM at 160 min, **Fig. 4E**), similarly to glucose-dependent growth (**Fig. 4C**). The fact that no significant differences were observed for this flux between glucose and gluconate cultures strongly supports our hypothesis that direct glucose uptake by *P*. *putida* is stably maintained at ∼7% if the sugar as well as gluconate are both present in the medium. The frequent sampling made it possible to catch a decrease in the Glk flux as the glucose concentration transitioned from ∼10 μM (i.e. similar to the *K*M of the glucose uptake system) to 0 μM. Indeed, this flux decreased to zero upon glucose depletion and remained as such at all later time-points.

Gluconate processing dominated sugar uptake in this experiment (**Fig. 4D**), with GnuK flux at 81-89%. Upon complete glucose depletion, gluconate was exclusively taken up (99% ± 3.8%) until its concentration decreased to 0.7 mM. The uptake of 2KG *via* KguK was low (4%-10%) while glucose was present in the culture medium. Once glucose was depleted, the KguK flux was negligible (0.6%) as long as gluconate was available. When the gluconate concentration dropped to 0.7 mM, KguK rapidly superseded GnuK as the predominant sugar processing flux. This Glk-GnuK-KguK flux transition was a rather revealing result, as this highly hierarchical and sequential uptake of the different carbon forms had never been described to this level of temporal resolution. The prevailing view was that the relative activities of the GnuK and KguK kinases determine the ratio of gluconate-to-2KG utilization. Similar to the hierarchy determined for the sugar uptake fluxes, three distinct growth phases were evidenced in the quantitative physiology data, which corresponded to growth until glucose depletion, followed by gluconate consumption and finally 2KG-dependent growth (**Fig. S2** in the Supplementary Material). Upon glucose consumption, the specific growth rate decreased from μ = 0.78 ± 0.08 h^−1^ to 0.38 ± 0.04 h^−1^ (**Fig. S2A** and **S2B**), while the gluconate consumption increased from 13.8 ± 0.6 mmol gCDW^−1^ h^−1^ to 19.4 ± 1.1 mmol gCDW^−1^ h^−1^ (**Fig. S2C** and **S2D**). This shift was similar to the change in the specific rate of glucose consumption observed in the previous growth phase at 4.8 ± 0.4 mmol gCDW^−1^ h^−1^ (**Fig. S2B**). The formation of 2KG was very similar during both phases (11.3 ± 0.4 mmol gCDW^−1^ h^−1^ and 12.7 ± 3.1 mmol gCDW^−1^ h^−1^, respectively). Hence, the total rate of carbon uptake in the early growth phase was 5.7 ± 0.8 mmol gCDW^−1^ h^−1^ and 2.5 ± 0.5 mmol gCDW^−1^ h^−1^ in the later phase. Interestingly, there was no flux through KguK at 343 min (**Fig. 4E**), when 2KG peaked at 7.8 mM with gluconate at less than half that concentration (3.8 mM, **Fig. 4D**).

Since the transition between the different carbon uptake regimes occurred swiftly, without the biphasic (diauxic) growth behavior typically associated to adaptive gene expression, we hypothesized that these changes could be associated to post-transcriptional regulation. As indicated above, the co-consumption of glucose and 2KG could be triggered by the complementary use of NADP^+^ and NADPH in both pathways—G6PDH produces NAD(P)H, whereas KguD consumes NADPH. Regulation of enzyme activity by NADPH availability is a common occurrence in bacteria [55,73] and likely to govern carbon uptake as well. Because gluconate metabolism is energetically favorable compared to that of 2KG (with an energy yield of ∼5.5 ATP *versus* ∼1 ATP until Pyr formation, respectively; **Fig. 2D**), catabolite repression may also play a role in controlling the hierarchical consumption of these oxidized substrates.

The changes in the peripheral carbon uptake routes on gluconate cultures were echoed by the EDEMP cycle fluxes (**Fig. 4E**). While gluconate and glucose were co-consumed, the Pgi flux remained at an average of 19.9% ± 2.5%, similarly to the previous experiments on glucose. In contrast, the Pgi flux was only 9.3% ± 0.4% when gluconate served as the sole carbon source, and gradually increased to ∼18% as 2KG uptake became predominant. Higher EDEMP cycling during substrate co-utilization may reflect insufficient flux capacity in the lower EMP pathway, from glyceraldehyde-3-phosphate to pyruvate—which in turn might be associated to efficient protein allocation [74]. This general argument is supported by the fact that the lower EMP reactions are less favored from a thermodynamic point of view and have higher protein costs than the reactions of the EDEMP cycle [8].

By combining the MFA data obtained through D-fluxomics with quantitative physiology parameters, we could calculate the exchange fluxes between glucose, gluconate and 2KG in the PGS (**Fig. 4F**). The transition from Glk to GnuK-KguK–predominant regimes when *P*. *putida* KT2440 grows on glucose was clearly captured by the distribution of fluxes in the PGS—at a degree of resolution that had been out of the scope of ^13^C-MFA so far. The next step was investigating how tightly coupled the oxidation of glucose and gluconate is to biomass formation, and we set to conduct further experiments to elucidate this potential correlation.

#### Growth rate-dependent accumulation of gluconate and 2-ketogluconate in P. putida

Several studies reported on the accumulation and secretion of gluconate and 2KG when *Pseudomonas* species and related bacterial species face iron, phosphate or magnesium starvation [23,75,76]. The general paradigm is that glucose oxidation routes are upregulated to help solubilizing nutrients. However, we consistently observed gluconate and 2KG accumulation by engineered *Pseudomonas* strains, e.g. C2 auxotrophs [45] and anthranilate producers [77], under nutrient-sufficient growth conditions. A feature common to all these setups, however, is that bacterial growth was reduced as the concentration of oxidized sugars increased. On this background, we questioned whether the trigger for organic acids accumulation by *P*. *putida* could be a reduction in the growth rate rather than a specific starvation stimulus. This hypothesis was further impelled by our observation that resting (i.e. non-growing) cells of *P. putida* rapidly oxidized glucose to gluconate. Decoupling glucose oxidation from growth could be connected, among other factors, to the need of dissipating the energy acquired in the oxidation steps if not otherwise used, e.g. by H^+^ leakage across the membrane [78], spontaneous ATP dephosphorylation [79] or water-forming NAD(P)H oxidation [80]. Therefore, a general mechanism was considered whereby carbon uptake is coupled to bacterial growth (i.e. at constant biomass yield), whereas the PGS activities are independent of growth. In this scenario, the accumulation of acids due to low growth rates would be merely an intrinsic characteristic of the system—rather than a consequence of differential genetic or metabolic regulation. To explore this possibility, we ran shaken-flask cultures of *P*. *putida* KT2440 on glucose where the growth rate was artificially lowered by supplementation of chloramphenicol, a bacteriostatic antibiotic, at levels below the minimal inhibitory concentration. Chloramphenicol inhibits protein synthesis [81], thereby slowing down bacterial growth. We periodically monitored bacterial growth, glucose consumption and formation of organic acids in these cultures until the primary carbon substrate was totally depleted (**Fig. 5A**). In the absence of any antibiotic, the specific growth rate of *P*. *putida* KT2440 cultivated in de Bont minimal medium supplemented with 20 mM glucose was μ = 0.66 ± 0.02 h^−1^—similar to our previous observations. When the *P*. *putida* cultures were added with chloramphenicol at 5 and 10 μg mL^−1^, the growth rates were reduced to μ = 0.46 ± 0.02 h^−1^ and 0.38 ± 0.01 h^−1^, respectively (**Fig. 5A**)—although the final OD600 was similar in all cultures. The glucose consumption profile was similarly protracted in the presence of chloramphenicol, with sugar depletion at 7-8 h regardless of the antibiotic concentration. Importantly, the cultures with reduced growth rates transiently accumulated the two oxidized forms of glucose at significantly higher concentrations than in control conditions, without antibiotic. Hence, these observations expose an inverse correlation of gluconate and 2KG synthesis and the growth rate of glucose-grown *P*. *putida*.

**Fig. 5.**
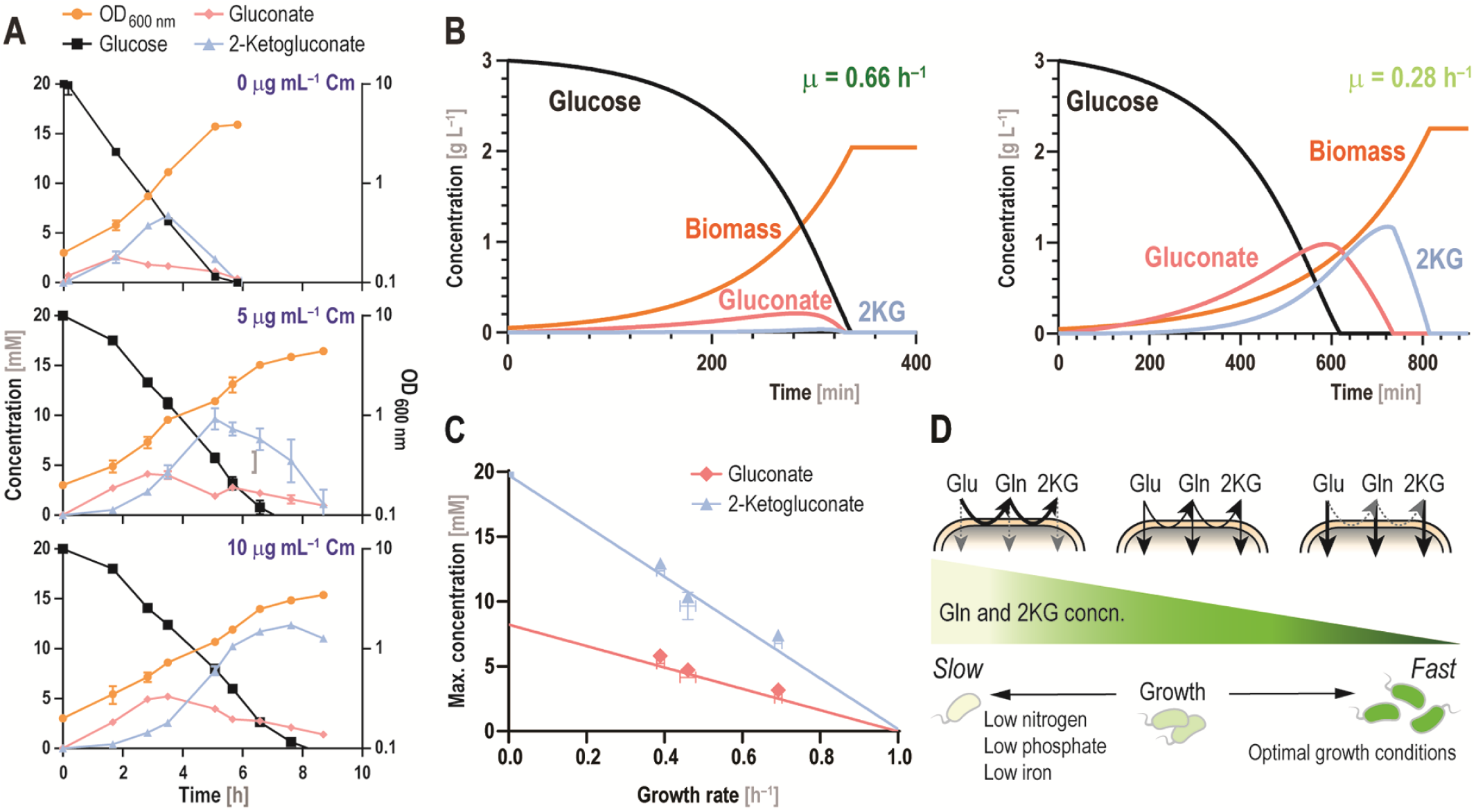
| Unveiling the mechanistic architecture of the periplasmic glucose shunt. **(A)** Growth profile and extracellular metabolite concentration for *P. putida* KT2440 grown on glucose in the presence of 0, 5 and 10 μg mL^−1^ chloramphenicol (Cm). Data represent mean values ± standard deviations of three independent experiments. **(B)** Modelling glucose consumption and gluconate and 2-ketogluconate (2KG) formation and uptake during batch cultivation of *P*. *putida* on glucose. Simulations were run with a growth rate μ = 0.66 h^−1^ (*left panel*) or with μ = 0.28 h^−1^ (*right panel*); other parameters used for *in silico* simulations are listed in **Table S2** in the Supplementary Material. **(C)** Accumulation of gluconate and 2KG, expressed as the maximum (max.) concentration, in glucose-grown *P*. *putida* KT2440 as a function of the specific growth rate (μ). The initial glucose concentration was 20 mM; μ was artificially altered by Cm supplementation [see **(A)**]. The data point at μ = 0 h^−1^ for 2KG was calculated assuming 100% conversion; linear regressions for gluconate and 2KG concentration over time (*t*) are [gluconate] = –8.24×*t* + 8.22 (*R*^2^ = 0.96) and [2KG] = –19.57×*t* + 19.74 (*R*^2^ = 0.98), respectively. **(D)** Proposed mechanism for organic acid formation in slow-growing bacterial cultures. Assuming that the total carbon uptake is directly correlated to cell growth and that sugar oxidation is growth-independent, higher amounts of gluconate (Gln) and 2KG are secreted by slow-growing cells. Hence, oxidation of glucose (Glu) and gluconate is favored over carbon uptake during slow growth. Conversely, carbon uptake increases in fast-growing cells, while sugar oxidation remains unaffected.

To further test our hypothesis in a formal framework, we developed an *in silico* model to simulate *P. putida* batch cultivations. In this model, the activity of the Gcd and Gad oxidases depends solely on the cell density, whereas the uptake of glucose, gluconate and 2KG is correlated with both the biomass concentration and the specific growth rate. The model was parameterized with values for specific enzyme activity, glucose uptake rate, biomass yield and KM values of the different transport systems taken from the literature on *P. putida* and related species (**Table S2** in the Supplementary Material). Simulating glucose-dependent, batchwise growth of *P*. *putida* KT2440 at μ = 0.66 h^−1^ (as experimentally determined, **Fig. 5A**) and 0.28 h^−1^ (i.e. a 60% reduction) predicted a significantly higher accumulation of gluconate and 2KG under slow-growing conditions (**Fig. 5B**). Interestingly, the model predicted no substantial glucose oxidation above a threshold growth rate, as the rate of consumption for the two acids would be higher than their rate of formation. Although the simplified model presented here neglects differential gene expression and protein allocation, which certainly influence glucose catabolism, it could predict gluconate and 2KG accumulation during glucose-containing batch cultivations of *P. putida* with a high degree of accuracy. Our simulations support the notion that acids accumulation is solely driven by the interplay of growth-coupled and growth-uncoupled reaction rates in the PGS. Indeed, when the experimentally-determined peak concentrations of gluconate and 2KG were plotted against the growth rate of glucose-grown *P*. *putida* KT2440, an inverse correlation was evident (**Fig. 5C**). The *y*-axis intercept for the [2KG] *versus* μ plot (i.e. μ = 0 h^−1^) was 19.7 ± 1.8 mM—thus closely matching the initial glucose concentration. The *y*-axis intercept for the [gluconate] *versus* μ plot was 8.2 ± 0.8 mM. Hence, these results suggest that non-growing *P. putida* cells would transform all glucose into 2KG with a maximum transient gluconate concentration of ∼8 mM. Conversely, the x-axis intercept (i.e. the μ value at which no organic acids are produced from glucose) was around μ = 1 h^−1^ for both linear regressions (**Fig. 5C**)—mirroring the results obtained in the simulations.

Based on these results, we propose a general mechanistic model for the PGS activities in glucose-grown *P*. *putida* (**Fig. 5D**), whereby higher amounts of gluconate and 2KG are secreted by slow-growing cells. From a physiological perspective, sugar oxidation may seem wasteful at a first glance. Yet, this phenomenon likely arises as a trade-off between energy efficiency and tailoring environmental niches towards optimal growth—as secreted organic acids have solubilizing effects on some minerals [22,82,83]. Furthermore, the PGS is relevant when glucose is highly abundant and not a growth-limiting nutrient—as reflected by the high *K*M of Gcd = 4.91 mM [53]. This metabolic scenario calls for great care in the analysis and interpretation of microbial physiology data, whenever the secretion of organic acids correlates with environmental stimuli that also influence the growth rate. Besides its relevance in Nature, the PGS is a key feature of *P. putida* amenable to engineering towards sugar-based biotechnological applications [84]. In this case, the correlation between growth rate and acid production can be adopted as a proxy to predict fermentation parameters [85].

## Outlook

The PGS is a widespread metabolic motif in environmental bacteria that has eluded a quantitative, mechanistic explanation thus far. Early experimental evidence has shown that a major fraction of hexoses are not processed by direct phosphorylation but rather oxidized to gluconate and 2KG when *P. putida* is cultured in the presence of glucose [26,27,58]. Moreover, a large part of the oxidized sugar pool is not taken up, but accumulated in the supernatant—becoming a substrate available for competitors in natural environments. What could be the advantage of bearing such a pathway for soil organisms?

Firstly, the total mineralization of glucose requires a considerable protein (enzyme) investment. As pointed out by Flamholz et al. [8], there is a trade-off between enzymatic cost and energy gained from a given glycolytic pathway. This principle explains why the (energetically favorable) EMP pathway is not widespread, but many bacteria rely on the ED pathway instead [10]. This occurrence is especially relevant in environmental microorganisms, e.g. the vast majority of marine bacteria [11], cyanobacteria [9] and several non-model organisms [66]. The ED pathway plays also a crucial role in the human microbiome, as an *E. coli* strain deficient in this route cannot colonize the large intestine [21]. The importance of gluconate for bacterial ecosystems is supported by its role as a signaling molecule for toxin production [20]. Secondly, and besides the biochemical reasons behind the coexistence of glucose processing pathways, competition between species is an important evolutionary driving force [86]. Rather than adopting the absolute best biochemical strategy, bacteria have to outcompete other species in their environmental niches [87]. The evidence presented here indicates that gluconate and 2KG secretion offers a competitive advantage that also helps understanding the widespread presence of the ED pathway. The PGS does neither require additional transporters nor initial phosphorylation of the carbon source, as do many other sugar processing pathways, and aerobic bacteria relying on the ED pathway could ‘steal’ sugar substrates from microbes utilizing solely the EMP—as they cannot consume gluconate or 2KG. Also, the enzymatic cost of the PGS is rather low (i.e. only two periplasmic oxidases, Gcd and Gad, are involved in the cognate reactions). Furthermore, each of the oxidase steps gains one H^+^, a very efficient energy yield as compared to other glycolytic routes, and, overall, the PGS is exergonic—but another glycolytic route has to be still employed as this oxidative sequence does not yield any precursors for biomass formation, besides ATP.

Developing and applying D-fluxomics permitted to determine all convergent fluxes in the PGS of *P*. *putida* KT2440. To the best of our knowledge, this is a first-case example on the use of D as a functional tracer for MFA towards resolving glucose oxidation. Furthermore, time-course D-fluxomics revealed the highly dynamic nature of glucose metabolism in *P. putida*, where spatiotemporally segregated uptake of the different sugar forms highlighted a hierarchical glucose, gluconate and 2KG processing—an aspect that has been largely neglected in metabolic models of *Pseudomonas*. This is not a surprising occurrence, as the data gained form ^13^C-MFA is usually provide as an average of substrate intake [30]. The majority of data reported in the literature was obtained during mid-exponential growth (usually, at OD600 ∼0.5) in batch cultures [33–35]. At this point, there is already considerable accumulation of gluconate and 2KG in the medium—therefore *P*. *putida* is exposed to a mixture of glucose and these metabolites, rather than growing exclusively on glucose as it does at the beginning of the cultivation.

With all of these considerations in mind, reporting the exact time point of sampling, together with the concentration of glucose and its oxidized derivatives, is of utterly importance to compare MFA results from different experimental setups. We note that this call for action is not a mere academic side-note, but it has important implications for metabolic engineering and bioprocess development. Indeed, our study demonstrates that the accumulation of gluconate and 2KG can be solely explained through a reduction in the growth rates, which could inform, for instance, feeding policies in fed-batch cultivations [88]. Interestingly, the activity of the PGS in *P*. *putida* does not follow the same principles as the overflow metabolism typical of *E*. *coli* and yeast (i.e. the Crabtree effect)— where a reduction in the growth rate limits the formation of acetate or ethanol in a glucose-rich environment [74,89,90].

Finally, even when we focused on resolving the PGS fluxes, the analytical power of the D-fluxomics methodology could be upgraded if further fluxes are elucidated. Our D-fluxomics protocol can be easily combined with other methods and is fully compatible with the use of traditional isotopic tracers, e.g. [1-^13^C]-glucose. Quantitative measurements of D-enrichment in downstream metabolites, e.g. intermediates from the tricarboxylic acid cycle and the PP pathway [91], can be also included to increase resolution and coverage. In this sense, adopting D-based isotopic tracers enables either elucidation of fluxes that could not be resolved with ^13^C-substrates or it improves analytical accuracy. These practical aspects are exemplified in our study, where the investigation of multiple isotopologues and isotopomers of F6P allowed for the resolution of fluxes within the PP pathway, the EDEMP cycle and the direct conversion of G6P to F6P *via* Pgi—as each of these pathways leads to a characteristic labeling pattern of F6P. While these developments will expand the scope of this methodology in the near future, we argue that D-fluxomics can be adopted as a powerful strategy for MFA of non-conventional microbial hosts.

## Acknowledgements

ACKNOWLEDGMENTS

The financial support from The Novo Nordisk Foundation (NNF10CC1016517 and NNF18CC0033664) and from the European Union’s *Horizon2020* Research and Innovation Program under grant agreement No. 814418 (*SinFonia*) to **P.I.N.** is gratefully acknowledged. The responsibility of this article lies with the authors. The NNF and the European Union are not responsible for any use that may be made of the information contained therein.

## SUPPLEMENTARY MATERIAL

**Table S1** | Transitions and optimized parameters for metabolite detection by MS.

**Table S2** | Parameters used for simulations of growth and glucose, gluconate and 2-ketogluconate secretion.

**Fig. S1** | Quantitative physiology parameters for *P*. *putida* KT2440 cultivations on glucose.

**Fig. S2** | Quantitative physiology parameters for *P*. *putida* KT2440 cultivations on gluconate and glucose as co-substrates.

