## Supplementary Material for "Time-resolved, deuterium-based fluxomics uncovers the hierarchy and dynamics of sugar processing by *Pseudomonas putida*"

**Table S1.** Transitions and optimized parameters for metabolite detection by MS.

| Metabolite | Mass | Parent ion | Parent mass [m/z] | Product formula (Hill notation) | Product mass [m/z] | DP [V] | EP [V] | CE [V] | CXP [V] | RT [min] |
|---|---|---|---|---|---|---|---|---|---|---|
| **Glucose** | $m$+0 | $C_6H_{11}O_6^-$ | 179 | $C_2H_3O_2^-$ | 59 | –36.2 | –10 | –23.1 | –5.8 | 0.94 |
| | $m$+1 | $C_6H_{11}O_6^-$ | 180 | $C_2H_3O_2^-$ | 59 | –36.2 | –10 | –23.1 | –5.8 | 0.94 |
| | $m$+2 | $C_6H_{11}O_6^-$ | 180 | $C_2H_3O_2^-$ | 60 | –36.2 | –10 | –23.1 | –5.8 | 0.94 |
| | $m$+6 | $C_6H_{11}O_6^-$ | 185 | $C_2H_3O_2^-$ | 61 | –36.2 | –10 | –23.1 | –5.8 | 0.94 |
| **6PG** | $m$+0 | $C_6H_{12}O_{10}P^-$ | 275 | $H_2O_4P^-$ | 97 | –56.4 | –10 | –22 | –5.24 | 13.8 |
| | $m$+1 | $C_6H_{12}O_{10}P^-$ | 276 | $H_2O_4P^-$ | 97 | –56.4 | –10 | –22 | –5.24 | 13.8 |
| | $m$+3 | $C_6H_{12}O_{10}P^-$ | 278 | $H_2O_4P^-$ | 97 | –56.4 | –10 | –22 | –5.24 | 13.8 |
| | $m$+6 | $C_6H_{12}O_{10}P^-$ | 281 | $H_2O_4P^-$ | 97 | –56.4 | –10 | –22 | –5.24 | 13.8 |
| **F6P** | $m$+0 | $C_6H_{12}O_9P^-$ | 259 | $C_3H_6O_6P^-$ | 169 | –36.4 | –10 | –16 | –9.7 | 6 |
| | $m$+1 | $C_6H_{12}O_9P^-$ | 260 | $C_3H_6O_6P^-$ | 169 | –36.4 | –10 | –16 | –9.7 | 6 |
| | $m$+1 | $C_6H_{12}O_9P^-$ | 260 | $C_3H_6O_6P^-$ | 170 | –36.4 | –10 | –16 | –9.7 | 6 |
| | $m$+3 | $C_6H_{12}O_9P^-$ | 262 | $C_3H_6O_6P^-$ | 169 | –36.4 | –10 | –16 | –9.7 | 6 |
| | $m$+3 | $C_6H_{12}O_9P^-$ | 262 | $C_3H_6O_6P^-$ | 170 | –36.4 | –10 | –16 | –9.7 | 6 |
| | $m$+3 | $C_6H_{12}O_9P^-$ | 262 | $C_3H_6O_6P^-$ | 171 | –36.4 | –10 | –16 | –9.7 | 6 |
| | $m$+3 | $C_6H_{12}O_9P^-$ | 262 | $C_3H_6O_6P^-$ | 172 | –36.4 | –10 | –16 | –9.7 | 6 |
| | $m$+6 | $C_6H_{12}O_9P^-$ | 265 | $C_3H_6O_6P^-$ | 172 | –36.4 | –10 | –16 | –9.7 | 6 |
| **G6P** | $m$+0 | $C_6H_{12}O_9P^-$ | 259 | $C_4H_8O_7P^-$ | 199 | –58 | –10 | –15.7 | –12 | 6 |
| | $m$+1 | $C_6H_{12}O_9P^-$ | 260 | $C_4H_8O_7P^-$ | 199 | –58 | –10 | –15.7 | –12 | 6 |
| | $m$+1 | $C_6H_{12}O_9P^-$ | 260 | $C_4H_8O_7P^-$ | 200 | –58 | –10 | –15.7 | –12 | 6 |
| | $m$+3 | $C_6H_{12}O_9P^-$ | 262 | $C_4H_8O_7P^-$ | 200 | –58 | –10 | –15.7 | –12 | 6 |
| | $m$+3 | $C_6H_{12}O_9P^-$ | 262 | $C_4H_8O_7P^-$ | 201 | –58 | –10 | –15.7 | –12 | 6 |
| | $m$+3 | $C_6H_{12}O_9P^-$ | 262 | $C_4H_8O_7P^-$ | 202 | –58 | –10 | –15.7 | –12 | 6 |
| | $m$+6 | $C_6H_{12}O_9P^-$ | 265 | $C_4H_8O_7P^-$ | 203 | –58 | –10 | –15.7 | –12 | 6 |
| **FBP** | $m$+0 | $C_6H_{14}O_{12}P_2$ | 339 | $O_3P$ | 79 | –37.8 | –10 | –73.6 | –6.5 | 15 |
| | $m$+1 | $C_6H_{14}O_{12}P_2$ | 340 | $O_3P$ | 79 | –37.8 | –10 | –73.6 | –6.5 | 15 |
| | $m$+3 | $C_6H_{14}O_{12}P_2$ | 342 | $O_3P$ | 79 | –37.8 | –10 | –73.6 | –6.5 | 15 |
| | $m$+6 | $C_6H_{14}O_{12}P_2$ | 345 | $O_3P$ | 79 | –37.8 | –10 | –73.6 | –6.5 | 15 |

*Abbreviations*: 6PG, 6-phosphogluconate; F6P, fructose-6-phosphate; G6P; glucose-6-phosphate; FBP; fructose-1,6-bisphosphate; DP, declustering potential; EP, entrance potential; CE, collision energy; CXP, collision cell exit potential; and RT, retention time.

**Table S2**. Parameters used for simulations of growth and glucose, gluconate and 2-ketogluconate secretion.

| Reaction | $K_M$ [mM] | Apparent $V_{max}$ [μM min$^{-1}$] | Specific activity [U mg$^{-1}$] | References |
|---|---|---|---|---|
| Gcd | 4.91 | 68 | 0.200 | [1,2] |
| GnuK | 0.12 | | 0.045 | [3,4] |
| KguK | 0.4 | | 0.007 | [3,5] |
| Glk | 0.009[a] | | 0.150 | [3,6] |
| Gad | 0.8[b] | | | [7] |
| Glucose uptake | 0.011[b]; 0.008[c]; 0.0003[d]; 0.0014[e]; 0.008[b]; 0.0083[a] | | 2.4† 0.030[e] | [6,8-11] |

Notes: † assuming 0.5 mg protein per 1 mg of cell dry weight (CDW).
[a] *P. putida* U; [b] *P. aeruginosa*; [c] *P. putida* CSV86; [d] *P. chlororaphis*; [e] *P. fluorescens*

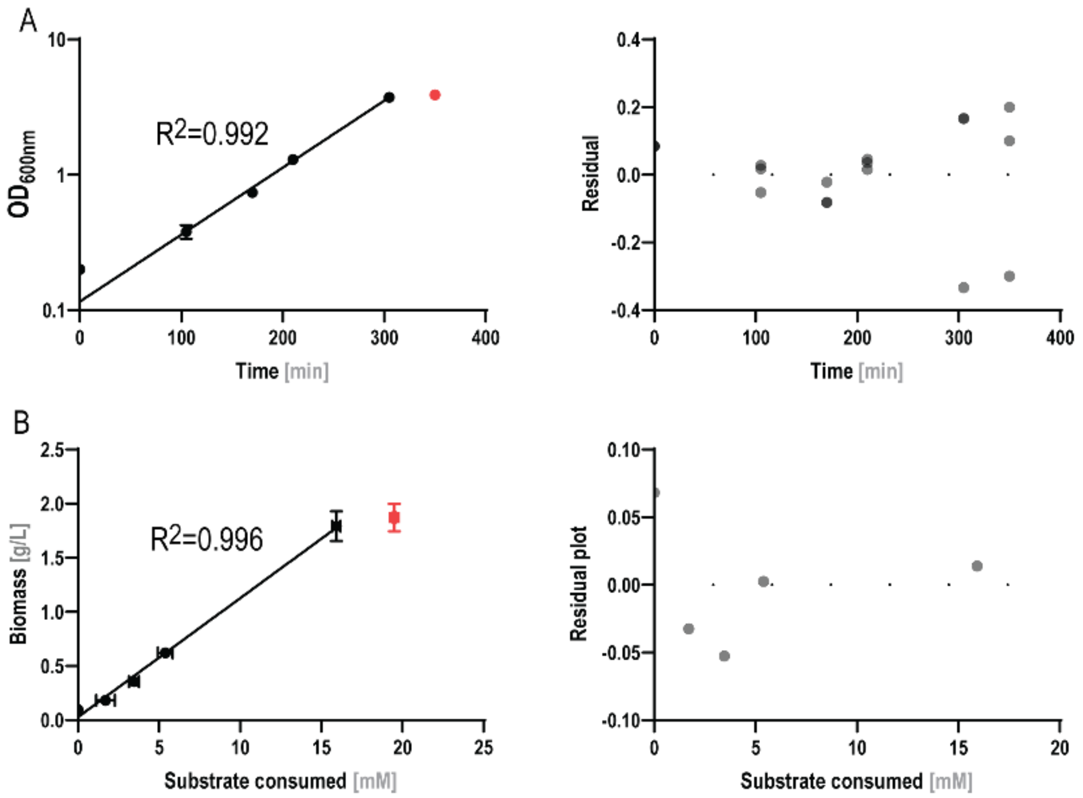


**Fig. S1 | Quantitative physiology parameters for *P. putida* KT2440 cultivations on glucose. (A)** Exponentially-fitted growth data. **(B)** Biomass yield on total substrate. Residual plots and $R^2$ are indicated for all fits; outliers are marked in red. $OD_{600\ nm}$, optical density measured at 600 nm. Results represent mean values ± standard deviations of three independent experiments.

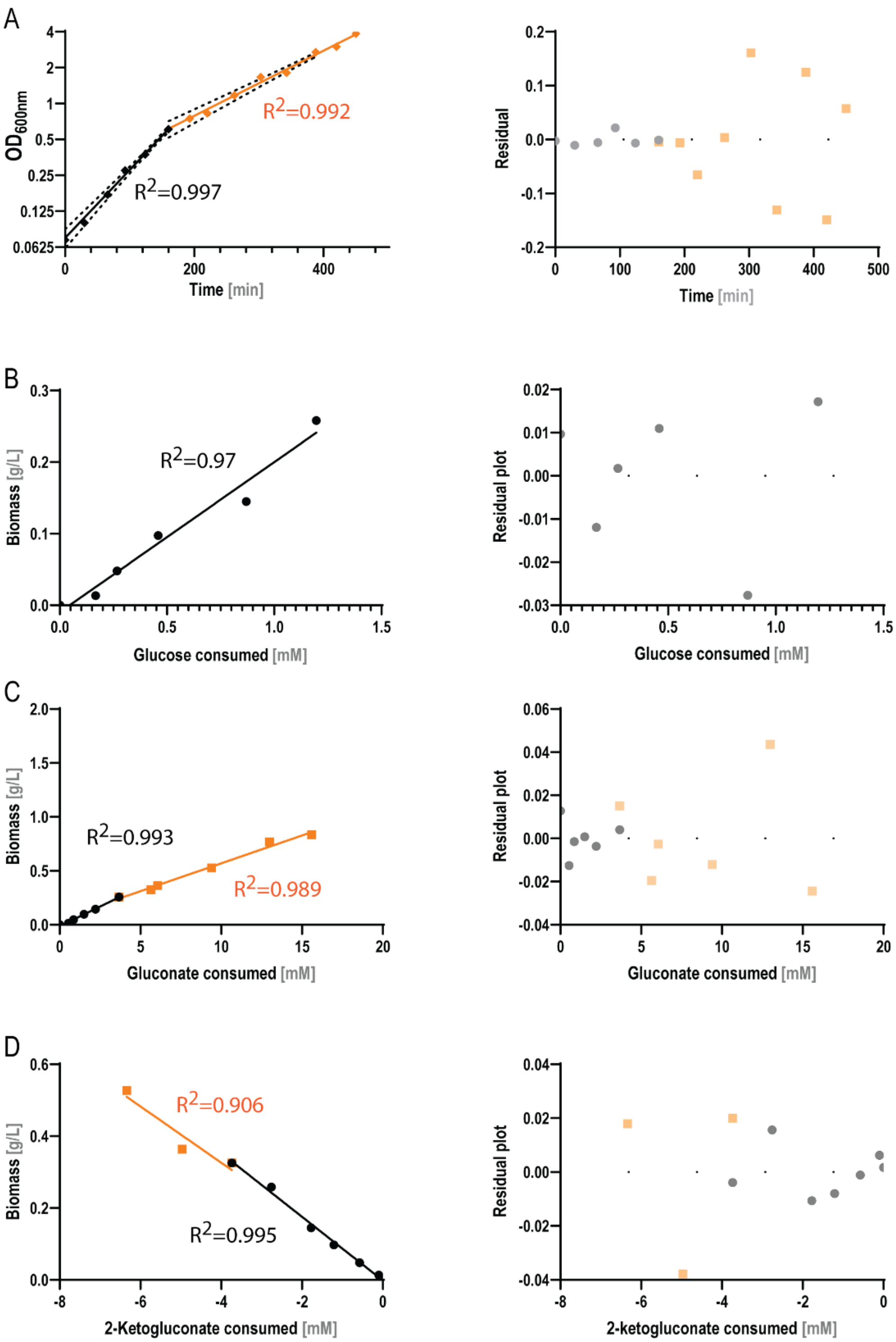


**Fig. S2 | Quantitative physiology parameters for *P. putida* KT2440 cultivations on gluconate and glucose as co-substrates. (A)** Exponentially-fitted growth data with (black symbols) and without (orange symbols) glucose present in the culture medium. The 95% confidence intervals are indicated with dashed lines. Biomass yields were calculated on glucose **(B)**, gluconate **(C)** and 2-ketogluconate **(D)**. Residual plots and $R^2$ are specified for all fits. $OD_{600\ nm}$, optical density measured at 600 nm. Results represent mean values ± standard deviations of three independent experiments.
